# Low Dose Hyperoxia Primes Airways for Fibrosis in Mice after Influenza A Infection

**DOI:** 10.1101/2020.06.17.157610

**Authors:** Andrew M. Dylag, Jeannie Haak, Rachel Warren, Min Yee, Gloria S. Pryhuber, Michael A. O’Reilly

## Abstract

It is well known that supplemental oxygen used to treat preterm infants in respiratory distress is associated with permanently disrupting lung development and the host response to influenza A virus (IAV). However, many infants who go home with normally functioning lungs are also at risk for hyperreactivity after a respiratory viral infection suggesting neonatal oxygen may have induced hidden molecular changes that may prime to the lung for disease. We discovered that thrombospondin-1 (TSP-1) is elevated in adult mice exposed to high-dose neonatal hyperoxia that is known to cause alveolar simplification and fibrotic lung disease following IAV infection. TSP-1 was also elevated in a new, low-dose hyperoxia mouse model (40% for 8 days; 40×8) that we recently reported causes a transient change in lung function that resolves by 8 weeks of age. Elevated TSP-1 was also identified in human autopsy samples of BPD-affected former preterm infants. Consistent with TSP-1 being a master TGFβ regulator, an early transient activation of TGFβ signaling, increased airway hyperreactivity, and peribronchial inflammation and fibrosis were seen when 40×8 mice were infected with IAV, which was not seen in infected room air controls. These findings reveal low dose of neonatal hyperoxia that does not affect lung function or structure may still change expression of genes, such as TSP-1, that may prime the lung for disease following respiratory viral infections, and may help explain why former preterm infants who have normal lung function are susceptible to airway obstruction and increased morbidity after viral infection.

## INTRODUCTION

It is well accepted that early oxygen (O_2_) exposure in preterm infants can disrupt lung development and function in extremely low gestational age newborns (ELGANs, <29 weeks gestation), often resulting in Bronchopulmonary Dysplasia (BPD). Cumulative O_2_ exposure strongly predicts BPD diagnoses and severity, but approximately 40% of ELGANs escape the BPD “label” because they are weaned off O_2_ or respiratory support by 36 weeks’ corrected age despite having significant supplemental O_2_ exposure (29, 60). Former ELGANs *with and without* BPD experience increased morbidity that can be linked to cumulative O_2_ exposure with increased health care utilization, symptomatic airway disease, and asthma medication use (18, 43, 57). Former ELGANs are especially vulnerable to respiratory viral infections with increased early childhood hospitalizations for respiratory illnesses (12, 24, 51) through poorly understood mechanisms. Infection-related lung injury results in airway remodeling and longer-term airway hyperreactivity (AHR) (26, 36, 46, 65), despite increased prescriptions for asthma-related medications (53, 54). The airway dysfunction is *not* bronchodilator responsive, distinguishing it from asthma (23). Thus, there is an urgent need to uncover novel mechanisms responsible for airway dysfunction and wheezing in former ELGANs.

Neonatal hyperoxia is one of the most commonly used exposures in animal models to perturb lung development and model BPD (10). Mice are born in the saccular stage of lung development where airways continue developing and alveolar structure is in its primitive stages, analogous to ELGANs (4). The dose and duration of hyperoxia matters when modeling neonatal O_2_ exposure in mice. For example, multiple studies, including several from our own laboratory, show that severe hyperoxia (≥ 60% O_2_ for ≥ 4 days) creates a BPD-like phenotype (10, 63) with alveolar simplification, airway remodeling, and viral susceptibility (14, 25, 35, 38, 40, 41, 44, 49, 67), even after a long period of room air recovery. These previous models, however, are limited because they often use O_2_ doses higher than those seen in real-world NICU settings and cause such profound alveolar simplification that it is difficult to discern physiological changes in the airway. This led our laboratory to develop a translational model of low dose chronic hyperoxia (40% O_2_ for 8 days; 40×8) which causes transiently increased airway resistance and decreased lung compliance with AHR and airway smooth muscle hypertrophy at 4 weeks (21), consistent with other studies (61). Interestingly, abnormal lung function, AHR, and smooth muscle hypertrophy all resolve at 8 weeks (21), where mice are morphologically and functionally “normal.” Taken together, our model of lower O_2_ exposures in preterm infants in modern NICUs shows changes in airway function without overt signs of alveolar simplification. We suggest this may more accurately replicate preterm infants with increased respiratory morbidity after leaving the NICU without a diagnosis of BPD.

Neonatal hyperoxia also has functional implications when adult mice are challenged with Influenza A viral (IAV) infection. Previous studies by our laboratory have shown that adult IAV infected who received higher dose neonatal hyperoxia (100% for 4 days, 100×4) at birth experience persistent inflammation and parenchymal fibrosis (14, 25, 35, 40, 41, 67). We previously showed that 100×4 hyperoxia depletes cardiomyocytes in the lung which has implications on pulmonary hypertension. Secondary analysis of that dataset suggests hyperoxia stimulates extracellular matrix thrombospondin 1 (TSP-1) and several members of the A Disintegrin and Metalloproteinase with Thrombospondin motifs (ADAMTS) family of proteinases which have TSP-1 like activity. Since TSP-1 upregulates TGFβ activity, we hypothesized that 40×8 adult mice would be primed for increased fibrosis after IAV infection. Herein, we show that when challenged with the HKx31 H3N2 Influenza A Virus (IAV) adult 40×8 mice develop peribronchial fibrosis after infection not observed in room air (RA) controls, that pathologically resolves over time, but persistently decreases lung compliance, thus creating a model of transient morbidity.

## METHODS

### Animal Exposures and Infection

All protocols were approved by the Institutional Animal Care and Use Committee of University of Rochester (Rochester, NY) and were consistent with The Association for Assessment and Accreditation of Laboratory Animal Care International policies (Frederick, MD). Litters of C57Bl/6J (Jackson Laboratory, Bar Harbor, ME) were placed into room air (RA) or 40% oxygen from post-natal day (PND) 0-8 as previously described (21). Nursing dams were rotated every 24-48 hours. After exposure, pups were allowed to mature until PND 56 under room air conditions where a subset of naïve mice were harvested for pulmonary function or qRT-PCR analysis. Infected mice were lightly anesthetized with ketamine/xylazine mixture and given 10^5^ plaque forming units (PFUs) influenza A (x31/H3N2) virus, which was grown and titered in Madin-Darby Canine Kidney (MCDK) cells as previously described (64). Mice were weighed every other day for two weeks after infection, then weekly thereafter.

### Bronchoalveolar Lavage

Bronchoalveolar lavage (BAL) was performed in a subset of animals at post-infection day (PID) 3, 7, 10, and 14 with 3 separate 1 mL aliquots of ice-cold phosphate-buffered saline (PBS, FisherScientific, Hampton, NH), as previously described (14). The first supernatant was collected for protein analysis and frozen at −80°C for further analysis.

### Cell differentiation

BAL Fluid (BALF) from all 3 aliquots were combined, then separated by centrifugation with removal of erythrocytes in ammonium chloride lysing solution (0.15 M NH_4_Cl, 10 mM NaHCO_3_, 1 mM EDTA). Total cell count was measured with a TC20 Automated Cell counter (Bio-Rad, Hercules, CA). BALF was then transferred onto slides with a cytological centrifuge (Shandon Cytospin 2, Runcorn, UK) and stained with a Hema 3 Stain Set (FisherScientific, Hampton, NH). Images of stained cells were taken with a Nikon E800 microscope (Nikon Instruments Inc., Melville, NY) using a SPOT RT3 Camera and SPOT Imaging Software (v5.2, Diagnostic Instruments, Inc., Sterling Heights, MI). At least 200 cells were counted per slide with ImageJ (NIH, Bethesda, MD). Macrophages/monocytes, neutrophils, and lymphocytes were individually enumerated by two separate investigators.

### Protein analysis

BALF was analyzed using a DuoSet ELISA kit for Mouse CCL2/JE/MCP-1 and TGF-β1 (R&D Systems, Minneapolis, MN) according to the manufacturer’s instructions using a SpectraMax M5 Microplate Reader (Molecular Devices, San Jose, CA) and Softmax Pro 6.4 (Molecular Devices). The detection range for this assay was 3.9-250 pg/mL. Additionally, latent TGF-β1 was activated by incubating samples with 1N HCl and neutralizing with 1.2 N NaOH/0.5 M HEPES, as per kit instructions, and another ELISA performed to measure immunoreactive TGF-β1.

### Pulmonary Function Testing

Naïve (8-10 weeks old) and IAV-infected (14 and 56 days post-infection) mice were anesthetized with a ketamine/xylazine mixture [100 mg/kg (Par Pharmaceutical, Chestnut Ridge, NY) and 20 mg/kg (Acorn, Inc., Lake Forest, IL), respectively], immobilized with pancuronium bromide (10 mg/kg, Sigma-Aldrich, St. Louis, MO), and ventilated (SCIREQ Inc., Montreal, Canada) with a tidal volume of 10 ml/kg, 150 breaths/min, PEEP of 3 cm H_2_O, and FIO_2_ of 21% as previously described (21). Respiratory system resistance (R_rs_), Newtonian airway resistance (R_N_), respiratory system compliance (C_rs_), Elastance (H), Tissue Damping (G), hysteresivity (η, eta) were measured in triplicate at both time points.

### Human Tissues

Donor lungs samples were provided through the federal United Network of Organ Sharing via National Disease Research Interchange (NDRI) and International Institute for Advancement of Medicine (IIAM) and entered into the NHLBI LungMAP Biorepository for Investigations of Diseases of the Lung (BRINDL) at the University of Rochester Medical Center overseen by the IRB as RSRB00047606, as previously described (5, 7). Lung tissue sections were uniformly obtained from the right lower lobe of 6 infants, 3 infants born prematurely (25, 26, and 28 gestational weeks) that died between 84 and 86 weeks post-menstrual age with BPD (2 ventilator dependent, died of respiratory failure, 1 with chronic lung disease, but not vent dependent died of accidental event) and 3 infants, each born full term and died at 74 to 100 weeks post-menstrual age of other non-pulmonary causes (n =L3 in each group). No acute viral infections were reported in the past medical history of any infant. Sections (5 µm) of formalin inflated, paraffin embedded RLL parenchymal lung tissue blocks were de-paraffinized, re-hydrated, and stained for Anti-Thrombospondin 1 (ab85762, abcam, Cambridge, UK) and DAPI Fluoromount-G (SouthernBiotech, Birmingham, AL).

### Immunohistochemistry

At PIDs 14 and 56, right lobes of lungs were snap-frozen for qRT-PCR, and left lobes perfused with 10% neutral buffered formalin (NBF, Fisher Scientific, Hampton, NH) at 25 mm/Hg, embedded in paraffin wax, and cut to 4 µm thick. Additional samples were taken after saline flush with 1x PBS, before perfusion with NBF. Lung slices were stained with Hematoxylin and Rubens Eosin-Phloxine (H&E; Biocare Medical, Concord, CA) and Gomori’s Trichrome (Richard-Allan Scientific, San Diego, CA) for collagen.

Fluorescent immunohistochemistry was performed with primary antibodies S100A4 (FSP-1) (1:1000, PA5-82322, ThermoFisher Scientific, Waltham, MA), Anti-Influenza A Virus Nucleoprotein (NP; NR-43899, BEI Resources, Manassas, VA) or Anti-Thrombospondin 1 (ab85762, abcam, Cambridge, UK), with secondary antibody AlexaFluor 594 (1:200, A21207, ThermoFisher Scientific) and DAPI Fluoromount-G counterstain to view activated macrophages. Stained images were taken with a Nikon E800 microscope (Nikon Instruments Inc., Melville, NY) using a SPOT RT3 Camera and SPOT Imaging Software (v5.2, Diagnostic Instruments, Inc., Sterling Heights, MI). Photographs were analyzed with ImageJ.

### Fibroblast and collagen staining

FSP1 and Sirius red staining were quantified using ImageJ. Fluorescent images of each airway were taken under red (FSPI, Sirius red) and blue (DAPI) channels. A threshold for each fluorescence was set, and the same used for all images taken. Airway perimeter was measured to ensure only small airways analyzed, then enlarged by 40 (Sirius red) and 80 µm (FSP1). Particle area (Sirius red) and number (FSP1) were measured, to determine their prevalence surrounding the airways.

### qRT-PCR

RNA was extracted from tissue with TRIzol Reagent (Invitrogen, Carlsbad, CA) as previously described (69). Complimentary DNA was run on a C1000 ThermoCycler (Bio-Rad) using a Maxima First Strand cDNA Synthesis Kit (ThermoScientific). Quantitative real-time PCR was performed using iQ SYBR Green Supermix (Bio-Rad) with CFX96 Real-Time System (Bio-Rad). Genes of interest where run on plates with *mGapdh* as housekeeping gene, and analyzed using the ΔΔC_T_ method (33). Three to four samples per treatment were run in duplicate on each plate. Primer sequences can be found in Table 1.

**Table 1.**
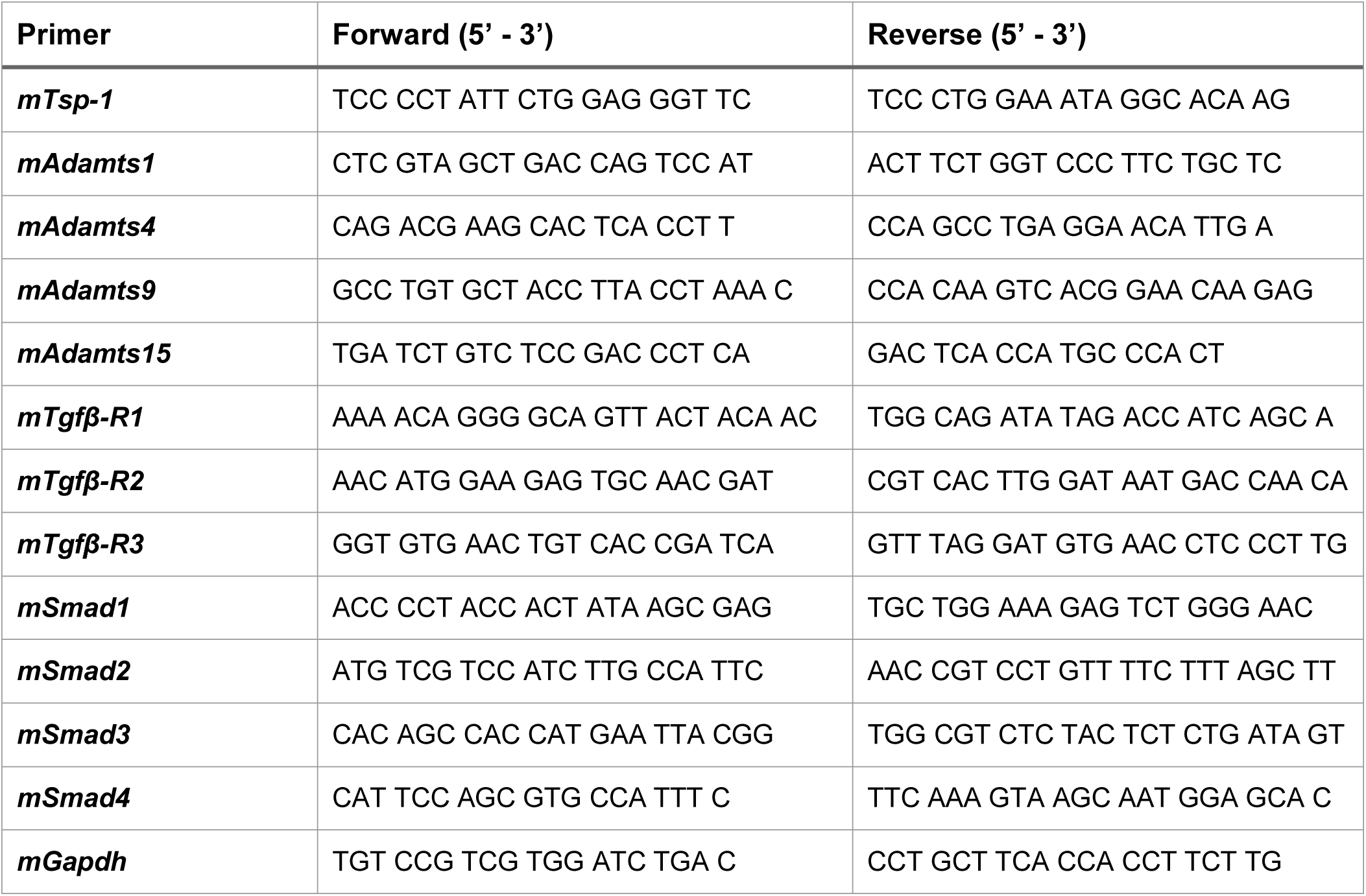
qRT-PCR Primer Sequences.

### Statistical Analysis

Statistical analyses were performed in GraphPad Prism (GraphPad Software v8, San Diego, CA). Pulmonary function data was subjected to D’Agostino & Pearson and Shapiro-Wilk tests for normality, Brown-Forsythe test for variance, and ordinary one-way ANOVA with Tukey’s multiple comparisons test for significance. In instances of failed normality or variance, Kruskal-Wallis non-parametic and Dunn’s multiple comparisons tests were performed for significance. Weight over time was tested for normality and variance, as described above, and multiple t-tests performed with Holm-Sidak correction. Cell differentiation data and ELISA data was subjected to Shapiro-Wilk test for normality. Holm-Sidak correction for multiple comparisons was used to further test cell differentiation and Kruskal-Wallis non-parametic and Dunn’s multiple comparisons tests were performed for significance on ELISA data. qRT-PCR data was analyzed using the ΔΔC_T_ method, and graphed as fold-change normalized to RA = 1. *P* values of ≤ 0.05 were considered significant for all analyses performed, and values graphed as mean ± SEM.

## Results

### Molecular differences persist in low-dose hyperoxia-exposed mice after Room Air Recovery

RA and 40×8 uninfected (naïve) mice were recovered in room air until 8-10 weeks of age (PND56, Figure 1A). We confirmed that RA and 40×8 animals have similar alveolar and airway structure by histology (Figure 1B), consistent with our laboratory’s previously published study (21). Furthermore, pulmonary function, measurements were similar between RA and 40×8 adult mice at this time point (Figure 1C). To determine whether adult RA and recovered 40×8 lungs had similar gene expression, we examined a previously published Affymetrix array in adult mice who received high-dose (100×4 oxygen at birth for candidate genes (68). Out of 45,109 probes present on the array, 54 transcripts were differentially expressed between the RA and the 100×4 mice using a false discovery rate of 10%. Neonatal hyperoxia reduced expression of 43 genes, most of which reflected a loss of pulmonary cardiomyocytes (68). We analyzed the upregulated transcripts for genes regulating inflammation using qRT-PCR and found the extracellular matrix protein thrombospondin 1 (TSP-1) and several members of the A Disintegrin and Metalloproteinase with Thrombospondin motifs (ADAMTS) family of proteinases were upregulated in the high dose 100×4 model (Figure 1D). TSP-1 and ADAMATS share common functions in their ability to activate latent TGFβ through conformational change of the latent binding protein that opens the binding site of TGFβ to its receptor (16). We confirmed that TSP-1 expression is increased 40×8 in adult mice and did not detect increased ADAMTS proteinases at high enough levels to justify further study (Figure 1D), thus choosing to focus on TSP-1.

**Figure 1.**
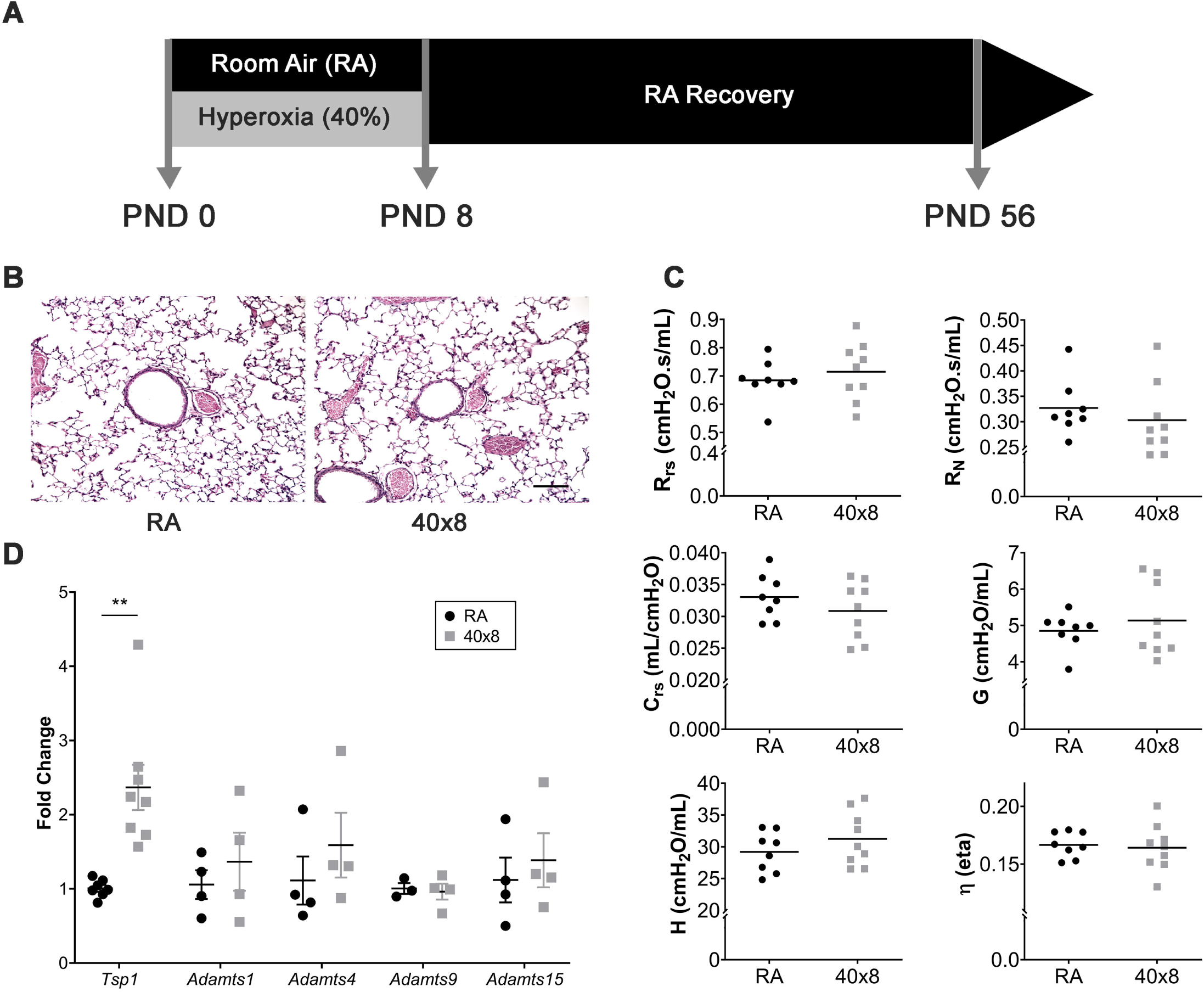
A) Model timeline of hyperoxia exposure and recovery in naïve mice. B) H&E sections of RA and hyperoxia (40% for 8 days; 40×8) exposed mice resemble RA controls at PND 56. C) Pulmonary function measurements are similar between RA and 40×8 mice for Respiratory System Resistance (R_rs_), Newtonian airway resistance (R_N_), Respiratory system compliance (C_rs_), Tissue Damping (G), Elastance (H), and hysteresivity (η, eta). *n* ≥ 8 per group. D) qRT-PCR of RA and 40×8 mice at PND 56 shows increased *Tsp1* and similar expression levels of other identified candidate genes. Data represent means ± SEM. ** *p* ≤ 0.01. Scale bar = 100 µm.

### TSP-1 expression is increased in hyperoxia-exposed mice and humans with BPD

To evaluate the prevalence and distribution of TSP-1, tissue sections of naïve 40×8 mice and BPD-affected human tissue samples were stained with TSP-1 antibody and counterstained with DAPI. In 40×8 mice, TSP-1 was increased in the alveolar spaces (Figure 2A) but less noticeable around the airways. In humans (N=3 controls, N=3 BPD lungs), increased TSP-1 was similarly detected in the alveolar spaces of 2/3 BPD infants where staining was much more sparse in control infants (Figure *2B*).

**Figure 2.**
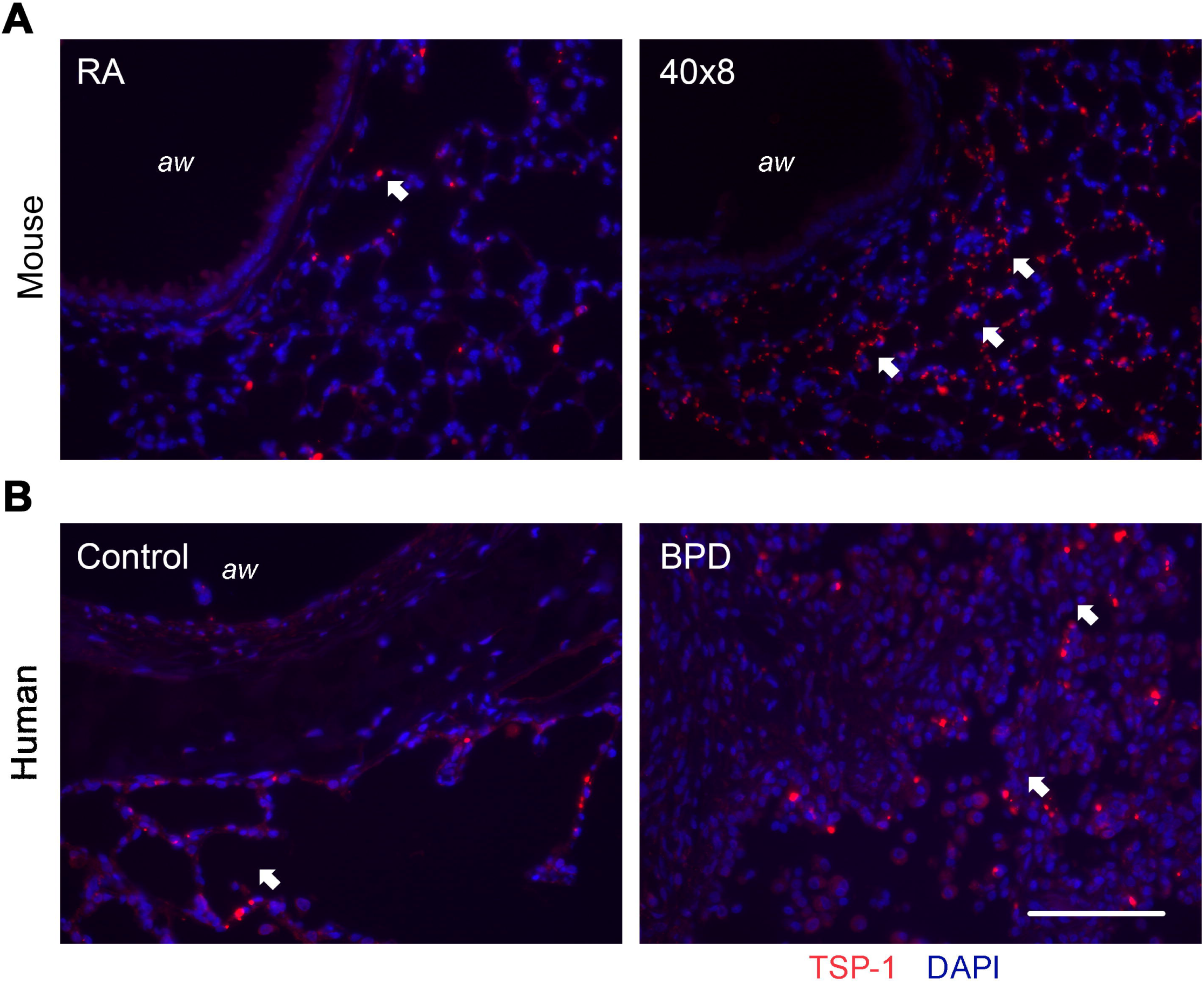
Lung samples were obtained from C57BL/6J A) mice at PND 56 that were exposed to oxygen at PND 0-8, and B) former premature infants at 1-2 year who passed away from BPD with non-BPD age-matched controls. Slides were stained with antibody to TSP-1 (red) and counterstained with DAPI (blue). White arrows indicate TSP-1+ cells. *aw* = airway. N = 3 samples per group. Scale bar = 100 µm.

### TGFβ signaling is altered in adult animals exposed to neonatal hyperoxia

To determine if key TGFβ signaling pathway mediators were different between RA and 40×8 animals, qRT-PCR was performed on naïve RA and 40×8 adult mice. We did not observe increased *expression of TGF*β-R1, or any of the canonical SMAD genes (Figures 3A, D, E, and F). Conversely, there was a strong trend for decreased *Tgfb-R2* and *Tgfb-R3* in naïve mice before infection (Figures 3B, C).

**Figure 3.**
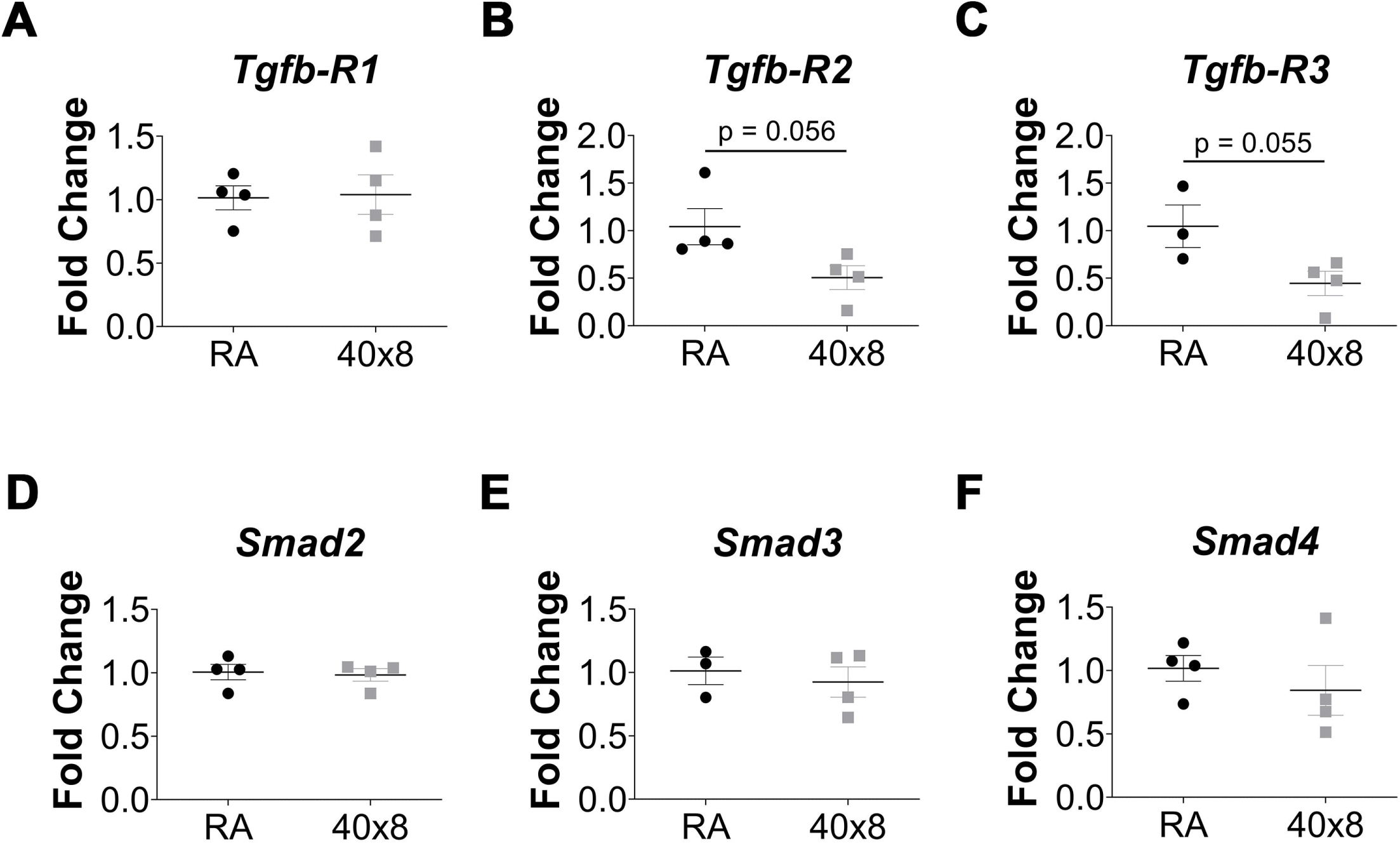
qRT-PCR was performed at PND 56 on naïve RA and 40×8 mice for A) TGFBRI, B) TGFBRII, C) TGFBR3, D) SMAD2, E) SMAD3, and F) SMAD4/co-smad. Data represent means ± SEM; *n* = 3-4 samples per group.

### Oxygen Exposed Mice Have Worse Airway Disease after Influenza Infection

Since TSP-1 activates TGFβ, we wanted to test the hypothesis that the 40×8 lung was primed to activate TGFβ signaling following an insult. We administered IAV to hyperoxia and RA exposed mice as a profibrotic challenge, because IAV causes fibrosis in mice and hyperoxia-exposed infants have increased morbidity after viral infections. To test this hypothesis, adult RA (RA-PBS or RA-x31) and 40×8 (O_2_-PBS or O_2_-x31) mice were nasally inoculated with HKx31 IAV or sham (PBS) (Figure 4A). Infection was confirmed by positive nucleoprotein (NP) staining in the airway club cells, indicating active IAV infection in both RA-x31 and O2-x31 animals (Figure 4B). Two weeks after infection, persistent airway inflammation (Figure 4C) and fibrosis (Figure 4D) were observed in O_2_-treated animals compared to RA controls. To confirm increased fibrosis in O_2_-x31 animals, Sirius Red collagen staining was performed (Figure 4E), imaged, and quantified in both groups with a focus on the peribronchial spaces. Again, we observed increased Sirius Red staining (Figure 4F) in 40×8 animals 2 weeks post-infection. Weight loss over the first two post-infection weeks were not different between the RA-x31 and O2-x31 groups during the first two post-infection weeks through 8 weeks post-infection, whereas sham animals appropriately did not lose weight (Figure 4G). There were no differences in weight loss between male and female mice (data not shown). The observed histologic changes resolved by 8 weeks post-infection (Figure 4H).

**Figure 4.**
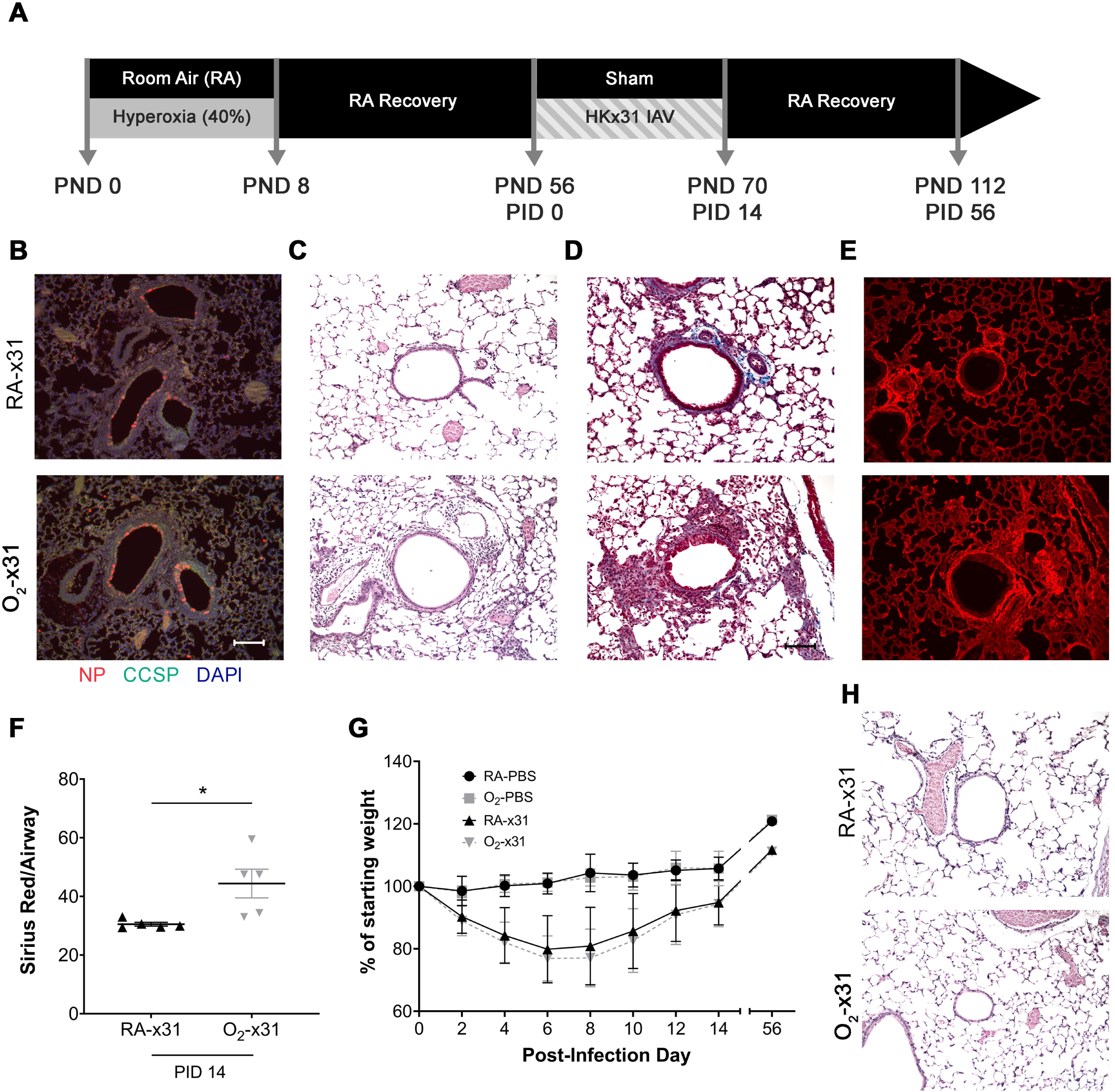
A) Experimental timeline for O_2_ exposure and IAV exposures. At PND 56, mice previously exposed to RA or 40×8 at PND 0-8, were intranasally infected with 10^5 PFU of H3N2 HKx31 IAV or PBS (sham). B) Fluorescent NP (red), counterstained with CCSP (green) and DAPI (blue), at PID3 indicates viral infection in small airways of both treatment groups. C) H&E and D) Trichrome stains at PID 14 indicate increased inflammation and fibrosis around the small airways of O_2_-x31 animals. E) Sirius red staining at 2 weeks post-infection shows increased collagen deposition in the O_2_-x31 treatment group. F) Increased Sirius red staining was detected around O_2_-x31 airways compared to RA. G) Animals of both groups that received IAV lost significantly more weight than sham mice, but similar weight loss occurred between infected groups. H) H&E staining at PID 56 showed a recovered phenotype with resolved inflammation and fibrosis for both groups. Data represent means ± SEM; *n* ≥ 5 samples per group. Scale bars = 100 µm. PND = post-natal day, PID = post-infection day.

Comprehensive assessments of pulmonary function were performed at both 2 and 8 weeks post infection to determine if the observed histologic findings had a physiologic correlate. At two weeks post infection, 40×8 animals had higher total Respiratory System Resistance (R_rs_, Figure 5A), Newtonian Airway Resistance (R_N_, Figure 5B), and Elastance (H, Figure 5E) with decreased Respiratory System Compliance (C_rs_, Figure 5C). Tissue damping (G, Figure 5D) and hysteresivity (eta, Figure 5F) were unchanged. The magnitude of the changes in R_N_ (which contributes to R_rs_) signify the majority of resistance change is due to changes in airway resistance. Interestingly, at 8 weeks post infection, the resistance changes normalized (Figures 5A-B), whereas the compliance and elastance (Figures 5C, 5E) remained persistently abnormal in the 40×8 animals where the RA animals returned to their previous levels. This suggests that lung function has reached a new, lower physiologic baseline, even after a long period of room-air recovery.

**Figure 5.**
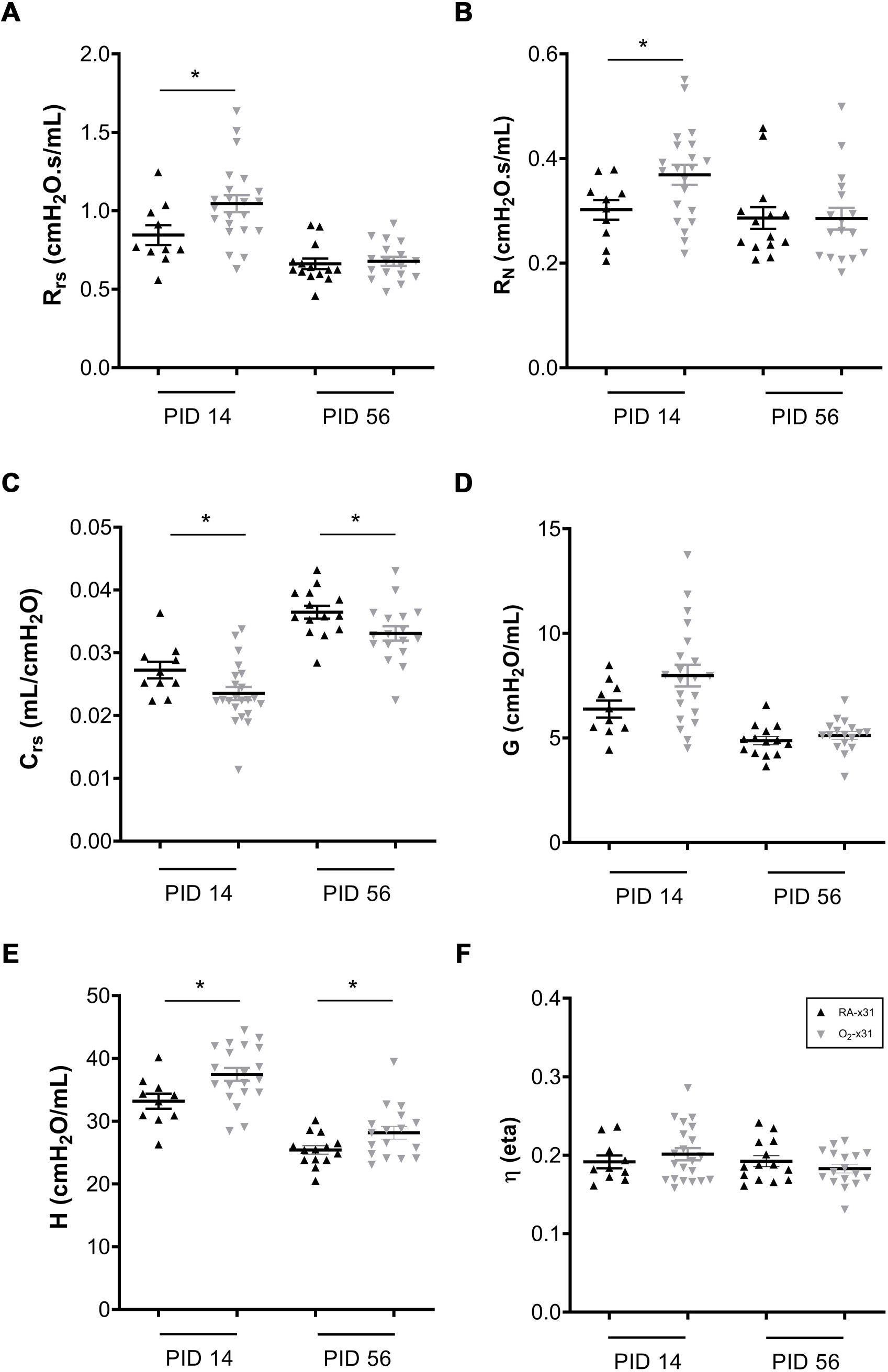
Pulmonary function testing at PID14 and 56. A) R_rs_ and B) R_N_ were higher in the O2-x31 group at PID14 indicating more hyperactive airways. C) C_rs_ was decreased in O_2_-x31 animals at both time points D) G was unchanged at both time points E) H was increased at both time points indicating increased tissue stiffness F) η remained unchanged at both time points indicating homogenous lung disease. Data represent means ± SEM; *n* ≥ 8 samples per group. * *p* ≤ 0.05. R_rs_ – respiratory system resistance, R_N_ – Newtonian resistance, C_rs_ – respiratory system compliance, G – tissue damping, H – tissue elastance, η (eta) - hysteresivity.

### Oxygen Exposed Mice have Persistent Peribronchial Inflammation

We next sought to determine whether low dose O_2_ exposure is associated with increased inflammatory cells similar to our previous studies using high-dose (100×4) oxygen (39). Bronchoalveolar Lavage Fluid (BALF) was obtained, spun, and quantified with differential cell counts at 4 time points during infection: days 3, 7, 10, and 14. Though we found a slight increase in total cell counts and neutrophils 7 days post infection in RA-x31 animals (Figure 6A-B), cell counts were otherwise similar between groups at the 3, 7, and 10 day time points. However, O_2_-x31 animals had trends for increased total and macrophage cell counts, 14 days after infection (Figure 6A, 6D). Quantification of MCP-1 protein was similar at all time points during infection (data not shown), distinguishing it from other studies in our laboratory (13), indicating that there may be alternate mechanisms driving increased inflammation in this model. To better characterize the inflammatory cells present around the airways, tissue sections were stained for fibroblast stimulatory protein 1 (FSP1), a marker shown to identify inflammatory subpopulations of macrophages (42). Increased FSP1 staining was concentrated around the small airways at 14 days post infection (Figure 6F) and was detected more frequently in O_2_-x31 animals (Figure 6G). Notably, FSP-1 staining was almost absent in the alveolar spaces with no differences detected between RA-x31 and O_2_-x31 animals (data not shown).

**Figure 6.**
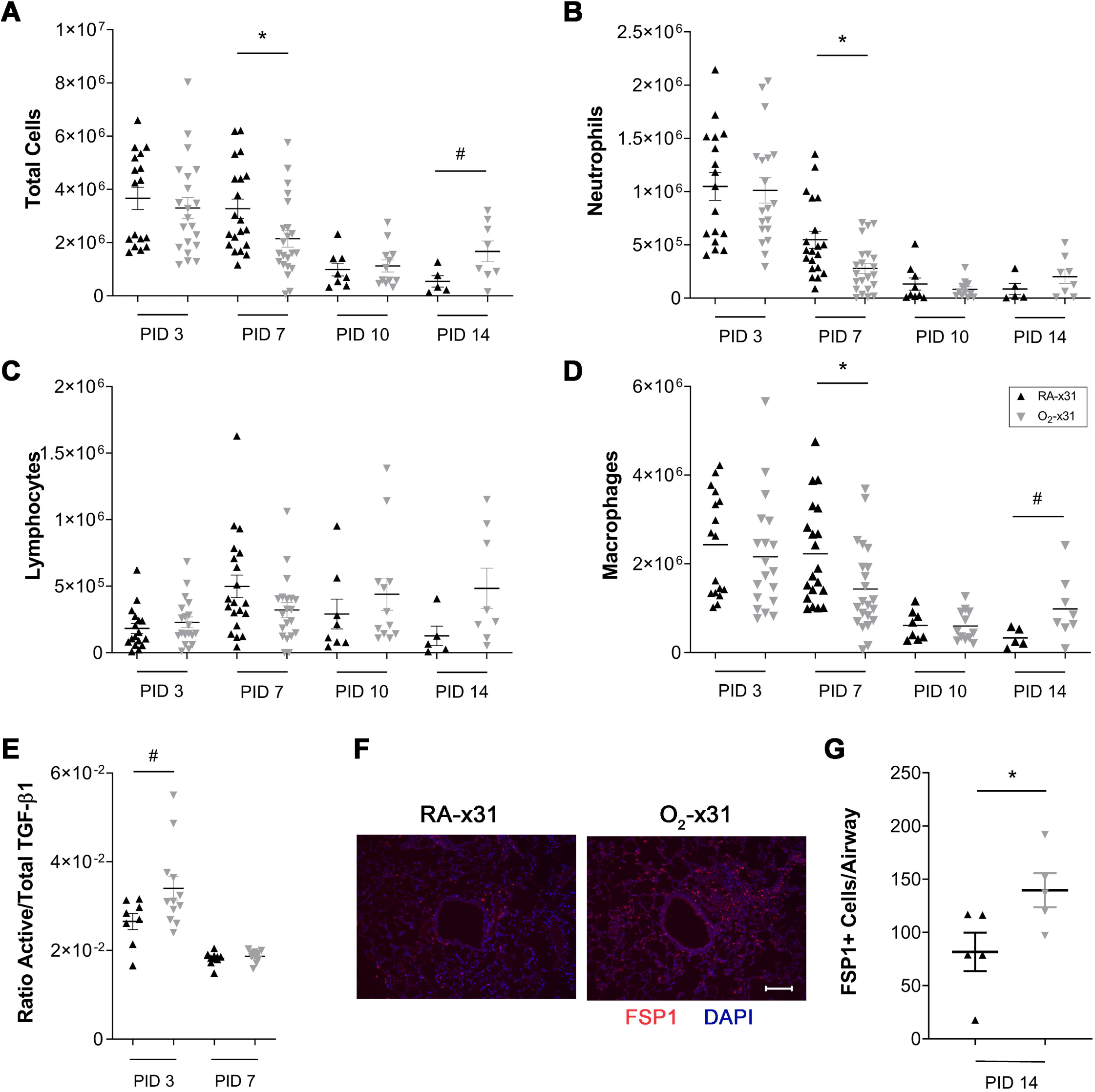
BALF was collected at PID 3, 7, 10, and 14 from RA-x31 and O_2_-x31 animals. A). Total cells, B) neutrophils C) lymphocytes, and D) macrophages were enumerated. Total cells were increased in the RA-x31 group at PID 7 and trends for increased total cells and macrophages were present at PID 14. E) ELISA was performed to determine ratio of activated/total TGF-β1 on PIDs 3 and 7 BALF. A trend towards a significant increase in O_2_-x31 protein at PID 3 was seen. PID 14 lung slices were stained with antibodies to FSP-1 (red) and DAPI (blue). F) More FSP-1+ cells were present around O_2_-x31 airways group compared to RA-x31 controls. Data represent means ± SEM; *n* ≥ 5 samples per group. # p < 0.10 (trend), * *p* ≤ 0.05, ** *p* ≤ 0.01.

The histologic, physiologic, and phenotypic differences in 40×8 animals led us to investigate possible mechanisms by which increased inflammation and fibrosis would be present and drive increased pulmonary morbidity after IAV infection. We analyzed BAL fluid for TGFβ using ELISA and observed a strong trend for increased active/total TGFβ 3 days post infection that normalized by 7 days (Figure 6E).

## DISCUSSION

Airway disease in former ELGANs is characterized by increased airflow obstruction in infancy (31), specifically in the mid-to-later forced expiratory flows (47, 52). These functional deficits predispose ELGANs to wheezing in infancy and childhood irrespective of their BPD status (8, 9, 15, 27, 28, 32, 56). Since BPD is often defined by need for supplemental O_2_ near term corrected gestational age (60), many infants who were exposed to O_2_ escape the BPD diagnosis, but still have significant pulmonary morbidity associated with their extreme prematurity. Our laboratory has performed several clinical studies quantifying cumulative O_2_ exposure in ELGANs *with and without BPD* and have shown that ELGANs with increased O_2_ exposure have worse obstructive lung disease (FEV_0.5_/FVC ratio), but “high” and “low” O_2_ exposed ELGANs have significant airflow obstruction (FEF_75_) compared to term infants at 1-year of age (22). Those functional studies were performed in asymptomatic, well ex-preterm infants, but there is an abundance of evidence that when challenged with a respiratory infection, former ELGANs have more significant lower airway symptoms with wheezing suggesting airflow obstruction and increased rehospitalization rates (20, 55, 65). These clinical studies emphasize that there is a spectrum of lung disease present in former ELGANs, and thus justifies studying a spectrum of O_2_ exposures in the laboratory that more closely model variant neonatal exposures, focusing on airway pathology to determine their impacts on lung development, function, and response to infection.

Using these ELGAN studies, we sought to create a translationally relevant paradigm of low-dose neonatal hyperoxia to show that mice are primed for airway disease when challenged with respiratory viral infection in adulthood. We chose 40% oxygen for 8 days because our previous studies showed that 40×8 mice have transiently increased airway hyperreactivity at 4 weeks of age that resolves by 8 weeks, such that the recovered animal is functionally and phenotypically indistinguishable from RA controls. This distinguishes it from other models wherein prolonged high O_2_ concentrations cause significant alveolar simplification, making it difficult to discern and isolate differences in airway pathology. Relatively low-dose 40×8 oxygen does not cause significant alveolar simplification, though if left in 40% oxygen for longer periods subtle changes in alveolar structure are detectable (37). We wanted to test if the “repaired” lung after hyperoxia would respond similarly to a RA animal when challenged with influenza A virus and hypothesized based on other studies from our laboratory (14, 25, 35, 40, 41, 67) that there would be persistently altered response to IAV, even after a long period of RA recovery. We chose HKx31 strain of influenza because it causes lower-airway symptoms, usually without mortality, and mice respond to IAV similarly compared to humans (19, 30, 58); unlike some other viruses (e.g. Respiratory Syncytial Virus) more commonly seen in infants that are difficult to model in mice. Our model is robust such that it recapitulates several observations in former ELGANs: 1) O_2_ causes changes in baseline airway mechanics, 2) airway changes are not overwhelmed by alveolar simplification, and 3) O_2_ exposed mice show increased severity of disease with viral infection. Additionally, many hyperoxia mouse models administer virus right when animals come out of hyperoxia, whereas this model allows for RA recovery and post-hyperoxia lung repair. Our 40×8 infection model occurs long after cessation of oxygen exposure, which has translational strengths to reflect former ELGANs later in infancy. Finally, other published models of airway dysfunction often treat animals with allergen or methacholine to observe differences in AHR. While this has been useful in identifying changes in airway smooth muscle, it may not explain airway disease of prematurity because these children often spontaneously wheeze following viral infections yet medications targeting airway smooth muscle relaxation (bronchodilators) are most often ineffective. Similarly, our infection-related changes are evident at baseline and without methacholine, implicating alternate mechanisms of airway pathology in O_2_ exposed mice apart from smooth muscle bronchospasm, distinguishing it from asthma.

We showed that 40×8 mice have increased airway-specific fibrotic repair resulting in hyperactive airways, associated with TGFβ hyperactivation, and delayed inflammatory resolution compared to room air controls. Our laboratory previously established a mouse model of high-dose neonatal hyperoxia (100% x 4 days; 100×4) followed by adult IAV infection associated with increased MCP1, marked *alveolar parenchymal fibrosis*, and increased mortality. In this paradigm, 100×4 mice have interstitial fibrosis, enhanced epithelial cell death, and increased mortality when exposed to IAV as adults (14, 25, 35, 40, 41, 67). These changes were partially attributed to the loss of type II alveolar epithelial cells (AEC2s) (69), but since 40×8 hyperoxia does not cause loss of AEC2s, other mechanisms driving disease severity were considered. IAV infection of 40×8 mice using the same virus increased <u>morbidity,</u> and was associated with peribronchial fibrosis 2 weeks after infection not observed in RA controls. Pulmonary function was abnormal with increased resistance and elastance and decreased compliance. We observed delayed clearance of immune cells 2 weeks after infection with increased staining for activated macrophages (fibroblast-specific protein 1, FSP-1 (42)) in 40×8 animals and trends for increased total and macrophage cell count in bronchoalveolar lavage fluid at that time point. IAV infection in 40×8 mice was not associated with increased weight loss or monocyte chemoattractant protein-1 (MCP-1) (13) (data not shown) as previously reported. Finally, 8 weeks after infection, the resistance changes resolve leaving behind subtle significant changes in compliance, establishing a model of intermittent morbidity that causes a downward shift in baseline lung function after infection in O_2_-x31 animals.

The peribronchial fibrosis changes in our model are associated with TGFβ hyperactivation. The TGFβ pathway regulates normal alveolar lung development (71). Global/floxed knockouts of TGFβ or key pathway mediators results in impaired alveolarization (50), but its role in airway development is more poorly defined. TGFβ1 signals through the canonical SMAD and/or non-SMAD dependent pathways to regulate gene expression, which in the lung can promote ECM collagen deposition and remodeling. The TGFβ signaling machinery changes both its expression and localization within the lung across developmental stages. Notably, TGFβ-R1 (receptor 1, ALK5), TGFβ-R2 (receptor 2), and SMAD3 decrease in expression throughout development, but change localization from vessels to airways as mice age such that when alveolarization is complete very little staining is evident in the alveolar spaces (3). This same study demonstrated that similar processes occur during human development, with most staining evident in the airways or vascular smooth muscle layer for ALK1, TGFβ-R1/ALK5, TGFβ-R2, and SMAD2. The evolutionary reason behind these changes throughout development have not been fully established, but may be concentrated in the areas of greatest lung growth during alveolarization. Dysregulation of the TGFβ pathway and its machinery has been implicated in hyperoxia-induced lung injury (2, 59, 66). Specifically, higher dose (85% O_2_ x 28 days) hyperoxia leads to a 4-fold increase and relocalization of the TGF-β2 receptor to the airway epithelium (2). Similarly, hyperoxia leads to a 6-fold increase co-SMAD/SMAD4 staining, also notable in the airway epithelium and alveolar septae (2). However, whole lung expression of these TGFβ pathway components in adult mice after 40×8 hyperoxia, was not increased, which led us to further investigate TSP-1 as a protein of interest.

Our finding of increased TSP-1 may provide mechanistic insight into IAV-related morbidity in hyperoxia exposed mice and humans with BPD. TSP-1 is a calcium binding ECM glycoprotein first discovered in platelet granules (6) and later localized to many other tissues including the lung (1). TSP-1 is synthesized by endothelial cells, fibroblasts, smooth muscle cells, monocytes, and macrophages (34) and interacts with several ECM components including integrins, fibronectin, cell receptors, growth factors (like TGFβ-1), cytokines, and proteases (11, 45). Antiangiogenesis, smooth muscle proliferation, nitric oxide signaling antagonism, and inflammation regulation are known functions of TSP-1 relevant to the lung (1). TSP-1 is required to form normal airway epithelium, as TSP1-null animals have bronchial epithelial hyperplasia, proximal mucous metaplasia, vascular smooth muscle hyperplasia, club cell hyperplasia, and uncontrolled inflammation (16). In contrast, upregulation of TSP-1 was noted in the preterm ventilated lung on autopsy (17), an in utero model of tracheal occlusion (62) (lung stretch), and other profibrotic diseases, but to our knowledge this has not been further explored in BPD. The BPD model is of particular interest because TSP-1 may play a role in capillary rarefication (also seen in mouse hyperoxia models (70)) and contribute to other hyperoxia-induced diseases such as pulmonary hypertension (48). Indeed, we confirmed increased alveolar staining for TSP-1 in both 40×8 exposed adult mice and BPD-affected human infants. The balanced regulation of TGFβ is vital for survival as TSP1-null and TGFβ-1-null mice both die of similar phenotypes (pneumonia) within weeks of birth (16). The strikingly similar phenotype of TSP1-null and TGFβ-1-null mice suggest that TSP-1 is the main TGFβ activator *in vivo* (16). Together, these previous studies on TSP-1/TGFβ create a potential “dual priming” effect of neonatal hyperoxia on the airway by the following mechanisms: 1) TGFβ receptor and signaling molecules localize to the airway throughout development and/or during hyperoxia exposure and 2) increased TSP-1 is primed to hyperactivate TGFβ. Thus, TSP-1 is an intriguing candidate for further study in rodent BPD models and in the extremely preterm infant.

Our results suggest low-dose neonatal O_2_ causes “silent” changes in gene expression with long-term functional consequences after a lung insult such as respiratory viral infection, and that the “repaired” lung still reacts abnormally to a profibrotic stimulus such as IAV. This 40×8 hyperoxia model causes an airway-specific phenotype observed at baseline and drives increased respiratory morbidity after infection, thus recapitulating airway diseases observed in former ELGANs. We can now exploit this model to better understand the origins of TGFβ hyperactivation (including TSP-1 as a candidate protein), and determine its source and specificity for the viral responses that may explain infection related morbidity in vulnerable former ELGANs.

## Acknowledgments

The authors would like to acknowledge Ethan David Cohen for his technical assistance and advice regarding the manuscript, and thank Biorepository for Investigation of Neonatal Diseases of the Lung (BRINDL) part of the Clinical and Translational Science Institute Informatics, Research Data Integration and Analytics group, University of Rochester Medical Center, for human tissue samples. We are extremely grateful to the families who have generously given precious gifts to support this research. We thank the members of the LungMAP Consortium for their collaborations and the members of the Pryhuber lab (Amanda Howell, Heidie Huyck, and Cory Poole) who prepared the human lung tissue.

## Author Contributions

AMD conceived the study and designed experiments. AMD and JH performed experiments with assistance from MY and RW. AMD and JH interpreted the data and wrote the manuscript. GSP curated the human tissue samples, and revised the manuscript. MOR conceived the study, revised the manuscript. All authors approved of the final version.

## Statement of Financial Support

Supported by a grant from the Strong Children’s Research Center at the University of Rochester and National Institutes of Health grant R01 HL091968 (MOR). NIH Center Grant P30 ES001247 supported the animal inhalation facility and the tissue-processing core. The human subject studies were supported by NHLBI Molecular Atlas of Lung Development Program Human Tissue Core grant U01HL122700 and U01HL148861 (G.H. Deutsch, T.J. Mariani, G.S. Pryhuber).

## Disclosure

The authors declare that no conflict of interest exists.

The authors have no financial conflicts to disclose and have agreed to submission of this article.

## Notes

### Competing Interest Statement

The authors have declared no competing interest.

